# Human cytomegalovirus infection changes the pattern of surface markers of small extracellular vesicles isolated from first trimester placental histocultures

**DOI:** 10.1101/2020.11.30.402693

**Authors:** Mathilde Bergamelli, Hélène Martin, Mélinda Bénard, Jérôme Ausseil, Jean-Michel Mansuy, Ilse Hurbain, Maïlys Mouysset, Marion Groussolles, Géraldine Cartron, Yann Tanguy le Gac, Nathalie Moinard, Elsa Suberbielle, Jacques Izopet, Charlotte Tscherning, Graça Raposo, Daniel Gonzalez-Dunia, Gisela D’Angelo, Cécile E. Malnou

## Abstract

Currently, research on the use of non-invasive biomarkers as diagnosis and prognosis tools during pathological pregnancies is in full development. Among these, placenta-derived small extracellular vesicles (sEVs) are considered as serious candidates, since their composition is modified during many pregnancy pathologies. Moreover, sEVs are found in maternal serum and can thus be easily purified from a simple blood sample. In this study, we describe the isolation of sEVs from a histoculture model of first trimester placental explants. Using bead-based multiplex cytometry and electron microscopy combined with biochemical approaches, we characterized these sEVs and defined their associated markers and ultrastructure. We next examined the consequences of infection by human cytomegalovirus on sEVs secretion and characteristics. We observed that infection led to increased levels of expression of several surface markers, without any impact on the secretion and integrity of sEVs. Our findings open the prospect for the identification of new predictive biomarkers for the severity and outcome of this congenital infection early during pregnancy, which are still sorely lacking.

## INTRODUCTION

Long considered as a passive barrier, the placenta is now recognized as a main actor in orchestrating the numerous exchanges between mother and fetus, in oxygen, nutrients and waste, protecting the fetus against infections and allowing adaptation of maternal metabolism to pregnancy [1, 2]. In the past decade, a new mode of communication of the placenta with both maternal and fetal sides has been described and extensively studied, consisting in the secretion of placental extracellular vesicles (EVs), which increases all along pregnancy and stops after delivery [3, 4]. EVs are membranous nanovesicles released by cells in the extracellular space and body fluids, under physiological and pathophysiological conditions [5, 6]. In a simplistic way, we can distinguish large microvesicles (up to 1 μm), derived from an outward budding of the plasma membrane; and exosomes, ranging from 30 to 200 nm of diameter, which are generated by inward budding of the membrane of late endosomes, leading to a multivesicular body that will fuse with the plasma membrane and release its content into the extracellular space. Discrimination between the different types of EVs based on their biogenesis pathway and/or physical characteristics is still the subject of intense work and numerous studies, and their classification is continuously evolving [5–8]. Hence, as it is often difficult to clearly prove the exact nature of exosomes compared to other vesicle subtypes, the term exosome has sometimes been used improperly in the literature, which must be interpreted with caution. We have therefore chosen in this manuscript to use the terminology small EVs (sEVs), according to the ISEV guidelines [9].

Interestingly, placental sEVS are detected in the maternal serum during pregnancy and their composition is altered upon placental pathologies, such as diabetes mellitus, intrauterine growth restriction or preeclampsia [10–16]. Thus, sEVs may represent valuable non-invasive biomarkers reflecting the status of the placenta and of the pregnancy [17, 18]. In order to identify such biomarkers for use in diagnosis or prognosis, it is important to develop relevant models which allow preparation of placental sEVs in a robust and reproducible manner, guaranteeing their purity for downstream analysis. In that matter, early placentas appear especially well suited experimental models, since many pregnancy pathologies and developmental defects are the result of placental insults occurring during the first trimester of pregnancy [19–21]. In this context, the use of tissue explants is particularly relevant, since they preserve the tissue cytoarchitecture, allowing to decipher the complex mechanism of (patho)physiological processes. Moreover, they also allow the study of the secretion of sEVs over several days, while this aspect limits the use of other currently available models [22, 23].

Among many environmental agents, viral congenital infections are a major cause of impaired placental and fetal development. Infection by human cytomegalovirus (hCMV) concerns 1% of live births in developed countries, and is responsible for various placental and fetal damages, especially at the level of the fetal central nervous system, leading to diverse brain disorders [24–26]. Non-invasive diagnostic tools to assess fetal hCMV infection are lacking and the gold standard diagnosis is using PCR on amniotic fluid, resulting in the necessity to perform an invasive amniocentesis, which is not devoid of risk [27]. Concerning prognosis, there are currently very few methods easily implementable to predict fetal impairment, especially concerning neurosensorial damage [28–30]. Thus, the identification of non-invasive diagnosis and prognosis biomarkers within sEVs would be a great step forward in assessing placental and fetal damage and would provide a valuable decision support tool.

In this context, we have adapted a histoculture model of first trimester placental explants that was developed in our team [31, 32], in order to isolate sEVs with a purity compatible with analyses of their composition and features. This model is permissive to hCMV infection [31, 32] and allowed us to purify sEVs devoid of contaminant viral particles. We showed that the secretion and integrity of sEVs was preserved upon hCMV infection, with significant modifications in the expression levels of some sEV surface proteins. Thus, this model opens up immense prospects for modeling chronic stresses at the start of pregnancy, like viral infections, to find biomarkers necessary to detect very early placental and fetal damage.

## MATERIALS AND METHODS

### Human ethic approval

The Germethèque biological resource center at the Toulouse site (BB-0033-00081) provided 21 placenta samples in order to carry out the research program. Except term of pregnancy, no other associated clinical data were collected, in accordance with policy concerning voluntary pregnancy termination. Germethèque obtained the agreement from each patient to use the samples (CPP.2.15.27). The steering committee gave its agreement for the realization of this study on Feb 5^th^, 2019. The biological resource center has a declaration DC-2014-2202 and an authorization AC-2015-2350. The hosting request made to Germethèque bears the number 20190201 and its contract is referenced under the number 19 155C.

### hCMV viral strain, viral stock production and titration

The viral strain of hCMV used in this study is the endotheliotropic VHL/E strain (a kind gift from C. Sinzger, University of Ulm, Germany) [33]. Viral stock was made by amplification of the virus on MRC5 cells and concentration by ultracentrifugation, as described previously [34]. Virus titration was determined by indirect immunofluorescence assays against the Immediate Early (IE) antigen of hCMV upon infection of MCR5 by serial dilutions of the viral stock [34]. Additionally, virus titration was also performed by qPCR as described on viral stocks and placental histoculture supernatants [35].

### Placental histoculture and infection

Placental histocultures were adapted from the model we previously described and validated (Figure 1 A) [31, 32, 36]. First trimester placentas (mean = 11.72 ± 0.39 (SEM) weeks of amenorrhea, *i.e.,* 9.72 ± 0.39 weeks of pregnancy) were collected following elective abortion by surgical aspiration at Paule de Viguier Maternity Hospital (Toulouse, France) by the medical team. Isolation of trophoblastic villi was performed from total placental tissue by manual dissection in Phosphate Buffer Saline (PBS), with particular care to exclude decidua, membranes and umbilical cord. Tissues were repeatedly washed in PBS to eliminate red blood cells. Each placenta was dissected in small pieces (2-3 mm^3^) and kept overnight in “Exofree” medium (see “Isolation of sEVs” section) in a 5 % CO_2_ incubator at 37°C, to eliminate the remaining red blood cells. To infect placental explants by hCMV upon dissection, the overnight incubation was performed with 500 μl of hCMV pure viral stock (corresponding to around 10^8^ ffu) mixed with 500 μl of Exofree medium. The day after (day 0), explants were washed six times in PBS and installed, nine by nine, on re-hydrated gelatin sponges (Gelfoam, Pfizer) in a 6-well plate containing 3 ml of Exofree medium per well (Figure 1 A). A minimum of six wells, *i.e.,* 54 explants, were used per experimental condition. Conditioned medium was collected and totally replaced with fresh Exofree medium every 3 to 4 days for the duration of the culture.

**Figure 1:**
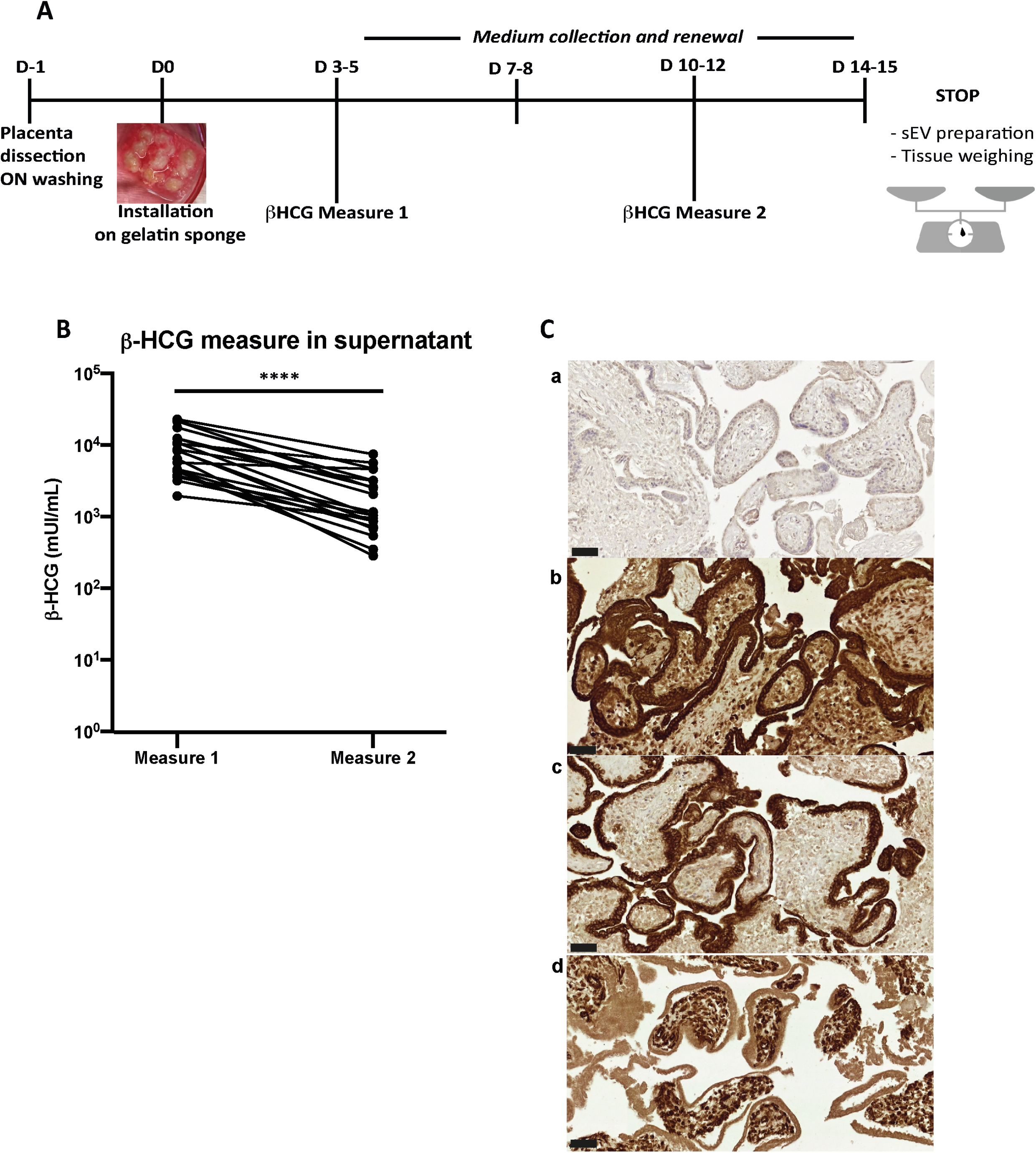
Placental histoculture set-up and characterization. A) Pipeline of placental histoculture model. B) β-HCG measurements in histoculture supernatant. For each placental histoculture, β-HCG measurement was realized in the supernatant between days 3-5 (measure 1) and days 10-12 (measure 2). **** *p<0.0001* by paired *t*-test (n=21 independent histocultures). C) Cross sections of immunohistochemistry and hematoxylin staining of placental villi from histoculture at day 14, observed by bright field microscope. a-Isotype control; b-Cytokeratin 7; c-PLAP; d-Vimentin. Image representative from at least three independent experiments. Scale bar = 50 μm.

To maximize the recovery of sEV and obtain a rate production compatible with further analyses, conditioned media were pooled and kept at 4°C until EV purification. At two time points during culture, 300 μl of culture supernatant were used to measure β-HCG levels. Free β-HCG was measured on a COBAS system (Roche Diagnostics, Switzerland), modular analytics E170, cobas e601 according to manufacturer protocol (Application Code Number 033) and according to a published method [37]. In addition to β-HCG dosage, release of virus by infected explants, indicating active viral replication, was assessed by hCMV qPCR titration on supernatant, as described above [35].

At the end-point of the histoculture, total collected medium was used to perform sEV isolation. Placental explants were weighted in order to normalize, calculate sEV yield and define an appropriate resuspension volume upon sEV preparation. Three explants were used for immuno-histochemistry and the others were frozen at −80°C for further analyses.

### Immuno-histochemistry

Placental explants were fixed in formalin during 24 h at room temperature and embedded in paraffin. Tissue sections (5 μm) were de-waxed using xylene and alcohol and epitope retrieval was carried out using citrate buffer (pH 6) at 95°C during 20 min. Sections were re-hydrated using TBS 0.01 % Tween 20 for 5 min and blocked with 2.5 % horse serum for 20 min. Immunostainings were performed with the following antibodies: rabbit anti-Cytokeratin-7 (Genetex; 2 μg/mL), mouse anti-Vimentin (Santa-Cruz; 2 μg/mL) and mouse anti-placental alkaline phosphatase (Biolegend; 1 μg/mL). Immunostaining for hCMV was performed as previously described [32], using a mouse monoclonal antibody directed against the hCMV IE antigen (clone CH160, Abcam). Secondary antibody-coupled to biotin was then used prior to Vectastain RTU elite ABC Reagent (Vector laboratories) and staining by diaminobenzidine (DAB). Sections were finally counter-stained with hematoxylin. Image acquisition was performed on a Leica DM4000B microscope or on a Panoramic 250 scanner (3DHISTECH).

### Isolation of sEVs

To purify sEVs from placental histocultures, culture media was depleted beforehand from EVs [9]. To this purpose, Dulbecco’s Modified Eagle Medium (DMEM with Glutamax, Gibco) supplemented with 20 % Fetal Bovine Serum (FBS, Sigma-Aldrich) was ultracentrifuged at 100,000 g for 16 hours at 4 °C (rotor SW32Ti, with maximal acceleration and brake) and filtered at 0.22 μm. “Exofree” medium was then obtained by a 1:1 dilution with DMEM to reach 10 % FBS and addition of antibiotics at the following concentrations: 100 U/ml penicillin - 100 μg/ml streptomycin (Gibco), 2,5 μg/ml amphotericin B (Gibco) and 100 μg/ml normocin (Invivogen).

All steps were then performed at 4 °C and PBS solution was filtered on a 0.22 μm filter. Procedures were adapted from [38–40] according to ISEV guidelines [9] and are presented in Figure 2A. From collected histoculture media, several differential centrifugation steps were carried out: a first preclearing centrifugation for 30 min at 1,200 g to eliminate dead cells and large debris, a second ultracentrifugation for 30 min at 12,000 g (rotor SW32Ti, with maximal acceleration and brake) to eliminate large EVs (principally microvesicles), and a last ultracentrifugation of the remaining supernatant for 1 hour at 100,000 g (Rotor SW32Ti, with maximal acceleration and brake) allowed to pellet sEVs. The pellet was then resuspended either in 100 μl PBS or in diluent C (Sigma) in order to stain the vesicles by the lipophilic dye PKH67 (Sigma) according to the manufacturer’s instructions (5 min incubation; 1:1,000 dilution). sEVs were then resuspended in a solution of 40 % iodixanol in sucrose and the last purification step was carried out by ultracentrifugation on a discontinuous iodixanol/sucrose gradient (10 to 40 % iodixanol) with deposition of the sEVs on the bottom of the tube, during 18 h at 100,000 g (rotor SW41Ti, acceleration 5, no brake). The fractions 2+3 of the six fractions harvested were then pooled and washed in 25 ml PBS. After a last ultracentrifugation for 1 h at 100,000 g (Rotor SW32Ti, with maximal acceleration and brake), the pellet was resuspended in PBS, in a volume proportional to the weight of tissue (1 μl PBS per 1 mg tissue). We submitted all relevant data of our experiments to the EV-TRACK knowledgebase (EV-TRACK ID: EV200049) and obtained an EV-METRIC score of 100 % [41].

**Figure 2:**
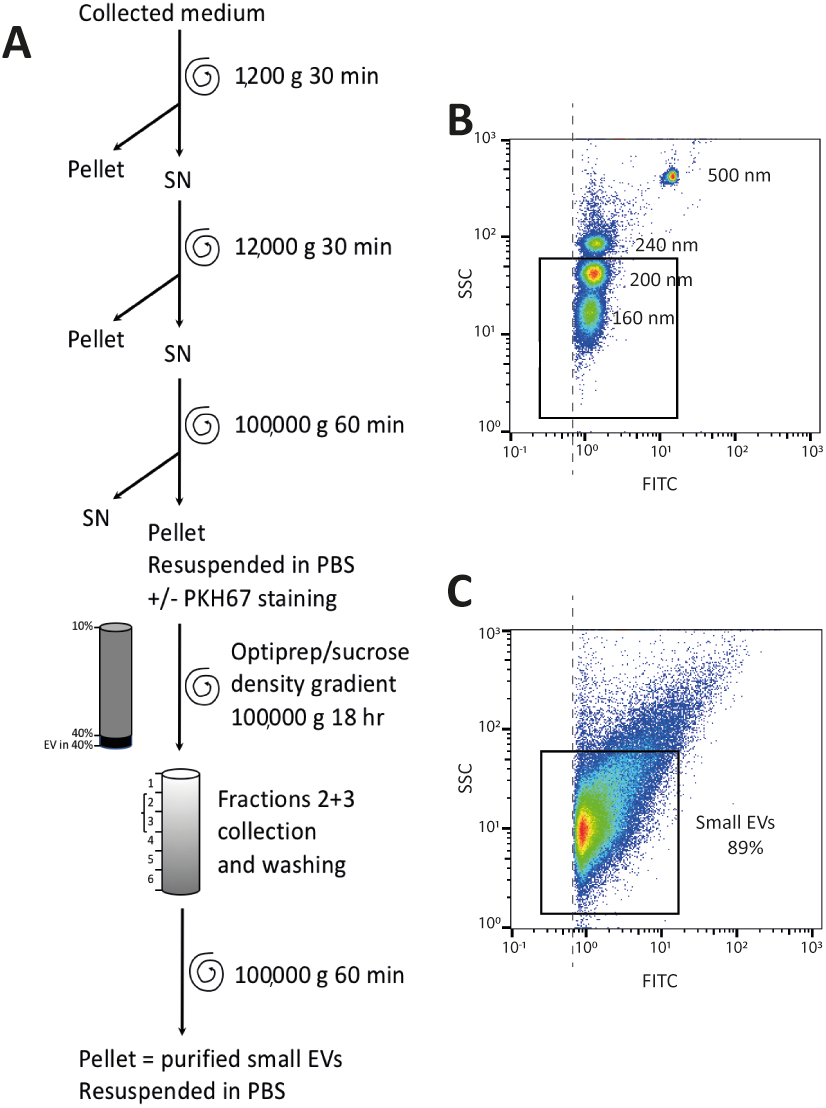
sEV preparation pipeline and flow cytometry analysis. A) Pipeline of sEV preparation using medium collected from placental histocultures. B) Flow cytometry standardization on fluorescent-FITC beads (160 nm, 200 nm, 240 nm and 500 nm). Each population size was defined on SSC granularity and FITC fluorescence parameters. The black rectangle indicates the gating strategy on small bead populations (160nm and 200nm). The dashed line represents the threshold for the detection of sEVs. C) Representative analysis of one placental sEV preparation gated on small events as described above. In this example, 89% of events were below 200nm.

### sEV flow cytometry

A Mascquant VYB Flow Cytometer (Myltenyi Biotec) was calibrated using Megamix-plus SSC FITC (Biocytex Stago) beads to standardize sEV measurements. Megamix-plus SSC beads of variable diameters (160 nm, 200 nm, 240 nm and 500 nm) were separated depending on size using SSC side scatter. A gating strategy was defined on 160 nm and 200 nm beads populations to analyze events of size below 200 nm (Figure 2B).

sEV preparations, previously stained with PKH67 as described above, were diluted 1:200 in filtered PBS and analyzed with the same parameters as those used for calibration beads. Gating on events of size below 200 nm allowed count of sEVs and calculation of their concentration for each preparation (Figure 2C). Each sample was analyzed twice. Data were then analyzed with FlowJo software (BD).

### Nanoparticle tracking analysis

sEV preparations were diluted 1:100 in filtered PBS (0.2 μm) and tracked using a NanoSight LM10 (Marvern Pananalytical) equipped with a 405 nm laser. Videos were recorded three times during 60 s for each sample at constant temperature (22 °C) and analyzed with NTA Software 2.0 (Malvern instruments Ltd). Data were analyzed with Excel and GraphPad Prism (v8) softwares.

### Transmission electron microscopy and immunolabeling electron microscopy

Procedures were performed essentially as described [42, 43].

For transmission electron microscopy (TEM), sEV preparations were loaded on copper formvar/carbon coated grids (Ted Pella). Fixation was performed with 2 % paraformaldehyde in 0.1 M phosphate buffer (pH 7.4), followed by a second fixation with PBS 1 % glutaraldehyde in PBS. Samples were stained with 4 % uranyl acetate in methylcellulose.

For immunolabeling electron microscopy (IEM), sEV preparations were loaded on grids and fixed with 2 % paraformaldehyde in 0.1 M phosphate buffer (pH 7.4). Immunodetection was performed with a mouse anti-human CD63 primary antibody (Abcam ab23792). Secondary incubation was next performed with a rabbit anti mouse Fc fragment (Dako Agilent Z0412). Grids were incubated with Protein A-Gold 10 nm (Cell Microscopy Center, Department of Cell Biology, Utrecht University). A second fixation step with 1 % glutaraldehyde in PBS was performed. Grids were stained with uranyl acetate in methylcellulose.

All samples were examined with a Tecnai Spirit electron microscope (FEI, Eindhoven, The Netherlands), and digital acquisitions were made with a numeric 4k CCD camera (Quemesa, Olympus, Münster, Germany). Images were analysed with iTEM software (EMSIS) and statistical studies were done with Prism-GraphPad Prism software (v8).

### Multiplex bead-based flow cytometry assay

sEV preparations were subjected to bead-based multiplex EV analysis by flow cytometry using the MACSPlex Exosome Kit, human (Miltenyi Biotec), according to the manufacturer’s instructions [44].

Briefly, sEV preparations were incubated overnight with 39 different bead populations, each coupled to a different capture antibody. The different bead populations are distinguishable by flow cytometry by a specific PE and FITC labeling. sEVs bound to the beads were then detected with a cocktail composed by anti-CD63, anti-CD9 and anti-CD81 antibodies coupled to APC. Beads coupled to isotype control antibodies were used to assess potential non-specific binding of sEVs. Background was also defined by performing the analysis without any sEVs. Flow cytometry analysis was performed with a MACSQuant Analyzer 10 flow cytometer (Miltenyi Biotec). The tool MACSQuantify was used to analyze flow cytometry data (v2.11.1746.19438). The background signals were subtracted from the signals obtained for beads incubated with sEVs. GraphPad Prism (v8) software was used to perform statistical analysis of the data.

### Western blot

sEV samples were lysed in non-reducing conditions in Laemmli buffer, heated for 5 min at 95 °C, and loaded on mini protean TGX precast 4-20 % gradient gels (Biorad) in Tris-glycine buffer for electrophoresis at 110 V for 2 h. Proteins were electro-transferred onto nitrocellulose membranes using the trans-blot turbo transfer system (Biorad) and membranes were blocked with Odyssey blocking buffer (Li-Cor Biosciences) for 1 h. Membranes were then incubated with primary antibodies: mouse anti-CD81 (200 ng/ml, Santa-Cruz), mouse anti-CD63 (500 ng/ml, BD Pharmingen) or mouse anti-CD9 (100 ng/ml, Millipore) overnight at 4 °C in Odyssey blocking buffer, followed by incubation with the secondary antibody IRDye 700 goat anti-mouse IgG (Li-Cor Biosciences), for 1 h at room temperature. Membranes were washed three times in TBS 0.1 % Tween 20 during 10 min after each incubation step and visualized using the Odyssey Infrared Imaging System (LI-COR Biosciences).

## RESULTS

To isolate sEVs and standardize their production from placental tissue, we adapted the placental histoculture protocol already developed and previously characterized by our team [31, 32]. To this aim, first trimester placentas were cultured as described in the Materials and Methods section and sampled at different time points (Figure 1A). To assess the viability of the placental explants [45], samples of the culture medium were used for β-HCG dosage, which revealed that placental explants secreted β-HCG at high levels (Figure 1B). In agreement with our previous studies [31], β-HCG levels gradually decreased, but remained sustained throughout the experiment. Assessment of tissue architecture and integrity was performed at the end of the culture and tissue sections were examined for the expression of Cytokeratin-7 (CK-7; trophoblast marker), placental alkaline phosphatase (PLAP; syncitiotrophoblast marker) and Vimentin (mesenchymal cell marker). We observed a typical double layer of trophoblastic cells, consisting of an outer syncitiotrophoblastic layer and an inner cytotrophoblastic layer, which surrounded the villous stroma (Figure 1C). Altogether, our results indicate that trophoblastic villi architecture was well preserved during the culture, as already shown in our previous works [31, 32].

We next designed a protocol to maximize the recovery of sEVs and allow their detailed characterization (Figure 2A). After the gradient ultracentrifugation step of this protocol, the majority of sEVs was found in fractions 2 and 3, corresponding to a density of, respectively, 1.086 and 1.116, consistent with the density expected for sEVs [8, 39]. The last sEV pellet was dissolved into a final volume of PBS proportional to tissue weight, to allow the normalization and comparison of sEV yields between experiments. Except when preparations were used for multiplex bead-based flow cytometry assays, sEVs were stained with the fluorescent lipophilic dye PKH67 before gradient ultracentrifugation, to allow their counting by flow cytometry (Figure 2A). Flow cytometer was calibrated with FITC-fluorescent beads of different sizes to define gating parameters before analysis of sEV preparations (Figure 2B). Only vesicles smaller than 200 nm of diameter were counted; the majority of the analyzed events displayed an approximate size smaller than 160 nm (Figure 2C). We obtained an average yield of 29,459 ± 5,370 sEV per mg of tissue (mean ± SEM).

To further describe the population of purified sEVs, we performed nanoparticle tracking analyses (NTA). Four independent sEV preparations were analyzed in triplicate. Concentrations of sEVs determined for the four preparations lied in the same range than the concentrations determined by flow cytometry and the comparison of the results obtained by the two methods showed no significant difference (Supplementary Table 1). The mode sizes for the four sEV preparations determined by NTA ranged between 125.7 and 170.7 nm, with a mean of 145.8 ± 9.3 nm (Figure 3).

**Figure 3:**
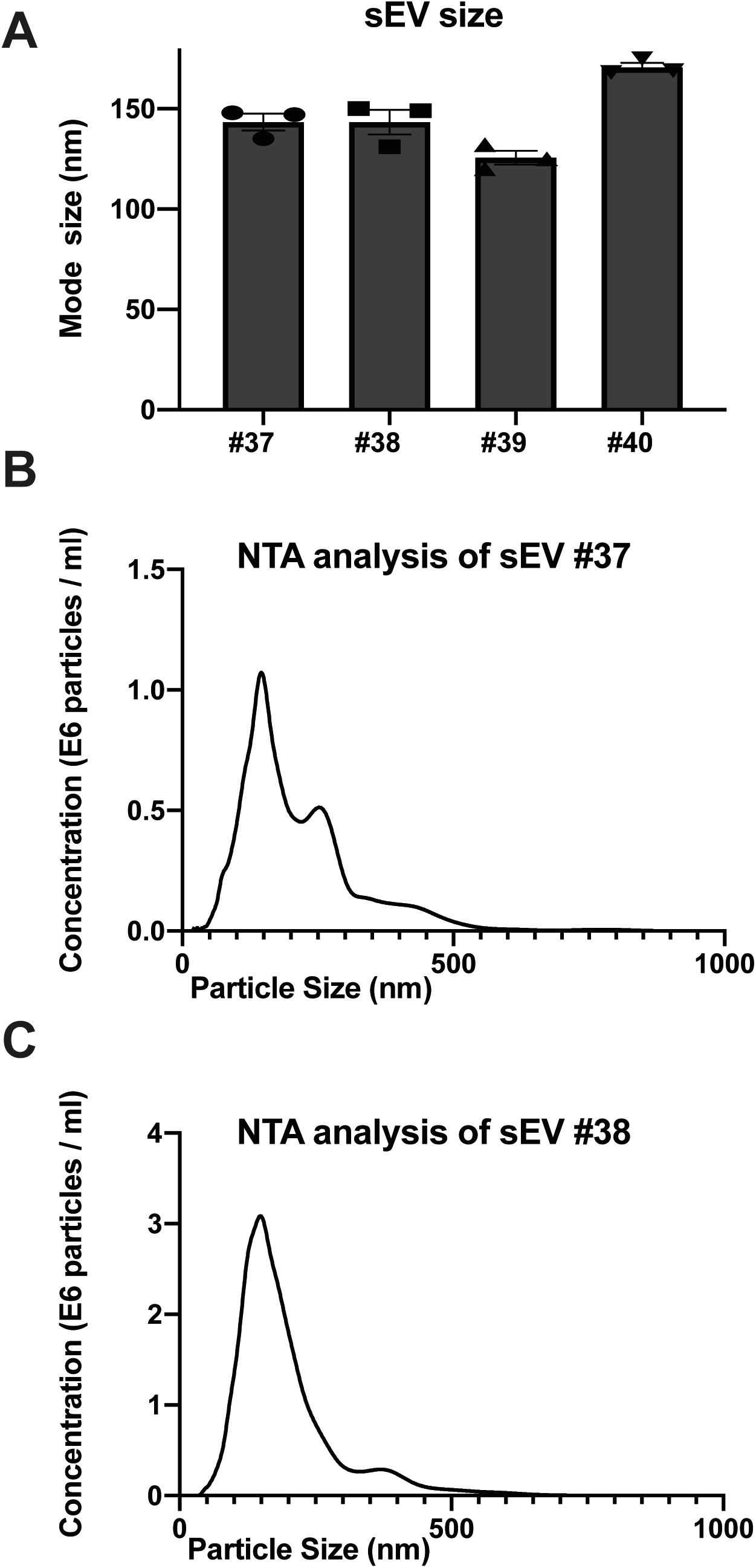
Nanoparticle tracking analysis of sEV prepared from placental explants. A) Individual analyses of mode size (nm) using four independent placental sEV preparations. Histograms represent mean ± SEM of three independent measurements (represented by individual dots) for each independent preparation. B) and C) Representative analyses of sEV size and concentration of two sEV samples (respectively, #37 in B and #38 in C).

Next, we performed an exhaustive morphological characterization of placental sEVs by TEM and IEM (Figure 4). The preparations were highly enriched in vesicles that presented the typical membranous appearance of sEVs (Supplementary Figure 1 and Figure 4A). The average relative size of sEV was determined using isolated sEVs from two independent sEV preparations. By focusing only on selected sEVs according to their structures, we measured an average diameter of 97 and 91 nm for the two preparations (Figure 4B). More precisely, by observing sEV size distribution, the majority of sEV diameters lied around 80 nm (Figure 4C). We next performed IEM to detect the canonical tetraspanin CD63, known to be enriched in endosome-derived exosomes. As observed in Figure 4D (and in Supplementary Figure 2), the majority of sEVs were highly positive for CD63. A manual counting of CD63-positive sEVs among total sEVs indicated that the percentage of CD63-positive sEVs was, respectively, of 60.32 and 61.83 % for the two sEV preparations.

**Figure 4:**
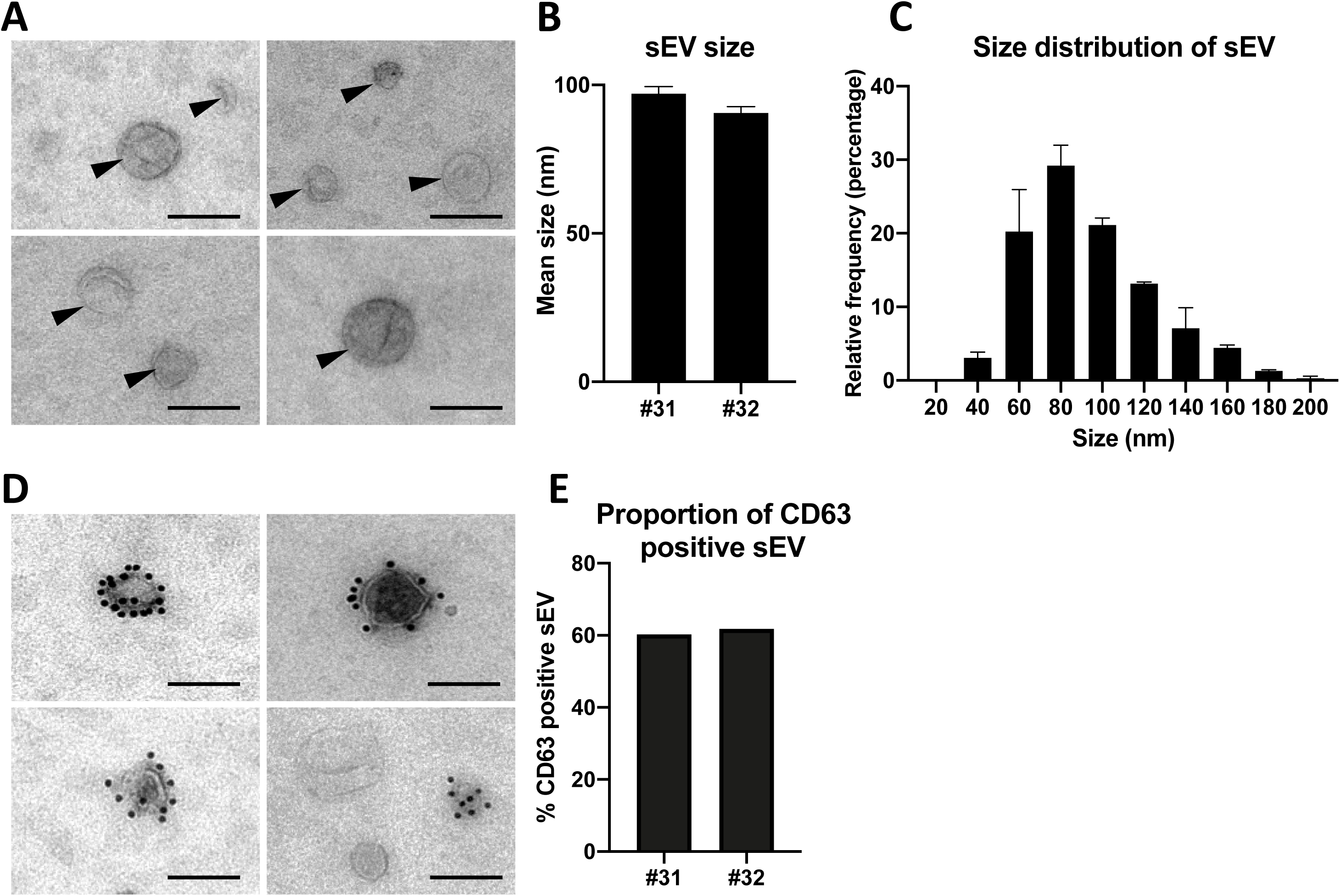
Electron and immuno-electron microscopy characterization of placental sEVs. A) Placental sEVs observed by TEM from two independent experiments. sEVs are indicated by black arrows. Scale bar = 100 nm. B) Mean size and min/max of placental sEVs for two independent experiments. Placental sEV size were measured manually with iTEM measure tool. C) Frequency distribution analysis of placental sEV size. Each bar of the histogram represents the mean ± SEM of the relative frequency per bin (bin width = 20 nm) of two independent experiments. Total sEV count was 172 for #31 and 208 for #32. D) Placental sEV were immunogold-labelled for CD63, revealed with Protein A-gold particle of 10 nm diameter and observed by TEM. Scale bar = 100 nm (n = 2). E) Percentage of placental sEV positive for CD63 (at least one bead counted per sEV) for two independent experiments. Total sEV count was 189 for #31 and 263 for #32.

Second, a multiplex bead-based flow cytometry assay was carried out to establish an exhaustive map of sEV surface markers (Figure 5). To this aim, we used an assay that allows for the simultaneous detection of up to 37 different EV surface markers in a semi-quantitative way [44, 46]. As shown in Figure 5A and expected from the IEM results, we observed a highly positive signal for CD63, a canonical exosome surface protein, which was also detected by western blot in two independent sEV preparations (Figure 5B). Two other canonical sEV surface proteins, CD9 and CD81, were found expressed in the sEV preparations (Figure 5A), but were not detected by western blot, probably because of the detection limit of the antibodies (data not shown). Altogether, these data indicate that isolated sEVs show the typical features of canonical exosomes, regarding ultrastructure, size and presence of typical exosome markers.

**Figure 5:**
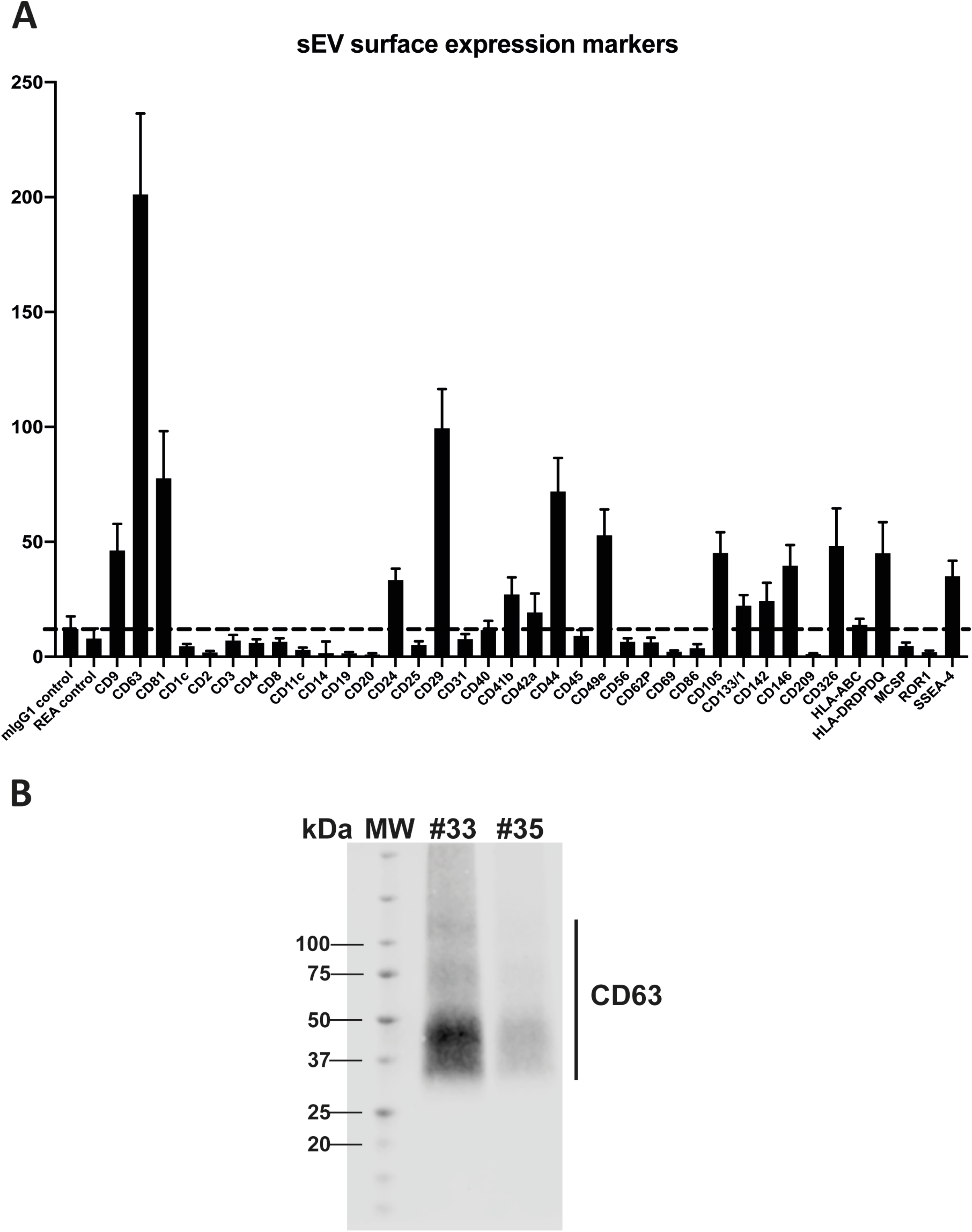
Placental sEV surface protein analysis. A) Surface expression level of several proteins of sEVs, based on the multiplex flow cytometry MACSPlex exosome kit assay. Each bar of the histogram represents the mean ± SEM calculated from 6 independent experiments, expressed in Median Fluorescence Intensity for different sEV markers indicated on the X axis. The dashed line represents the detection limit of the test (defined by the two first controls on the left of the histogram). B) Representative western blot analysis against CD63 performed on two independent sEV preparations (#33 and #35). CD63 appears as a smear, since the non-reducing conditions of the western blot preserve its rich glycosylated pattern. MW = molecular weight.

The bead-based flow cytometry assay also revealed that several proteins expressed by trophoblastic cells were present at the surface of sEV, including CD24 (which was already described on trophoblastic EVs) [47], CD49e (also known as integrin α5) [48], CD105 [49], CD146 (also named MCAM) [50] or CD236 (also known as EpCAM) [51]. Conversely, expression of non-trophoblastic markers such as CD4, CD8, CD31, CD45 or HLA-ABC [48, 52] was not detected on sEVs preparations. Of note, markers described for placental mesenchymal stem cells were also found at the surface of isolated sEV like CD29 (also known as integrin β1), CD44 and SSEA-4 [53, 54], indicating that these cells may also contribute to sEV secretion in the histocultures.

Next, we examined the impact of hCMV infection on the secretion and characteristics of placental sEVs. Placental explants were infected by the VHL/E clinical strain overnight, extensively washed and maintained in culture during two weeks to favor virus dissemination, as already described by our team [31]. To monitor virus release into the culture medium, we sampled one aliquot of histoculture supernatant after two medium renewal steps (corresponding to virus released between days 7 and 11), which was analyzed by qPCR. Placental explants displayed active viral release, with hCMV titers in the supernatant comprised between 1.05 10^4^ and 1.53 10^7^ copies/ml, the median lying at 3.04 10^5^ copies/ml (Figure 6A). Importantly, these titers were indeed due to virus release and did not correspond to remaining inoculum, since no viral genome could be detected by control qPCR experiments using UV-irradiated virus (data not shown). Moreover, analysis of the tissue sections by immunohistochemistry at the end of the culture confirmed the presence of the IE viral antigen (Figure 6B). Some cells showed intense staining, demonstrating that the virus had disseminated well into the tissue after two weeks. Viral infection did not modify the weight of tissue upon culture compared to non-infected conditions (Figure 6C), neither the level of secreted β-HCG that remained similar between non-infected and infected placentas for both measures 1 and 2 (Figure 6D). Finally, the tissue architecture remained well preserved, as attested by immunohistochemistry performed against CK-7, PLAP and Vimentin (Figure 6E), thereby guaranteeing that the sEV preparations were valuable and did not correspond to sEVs isolated from dying tissues.

**Figure 6:**
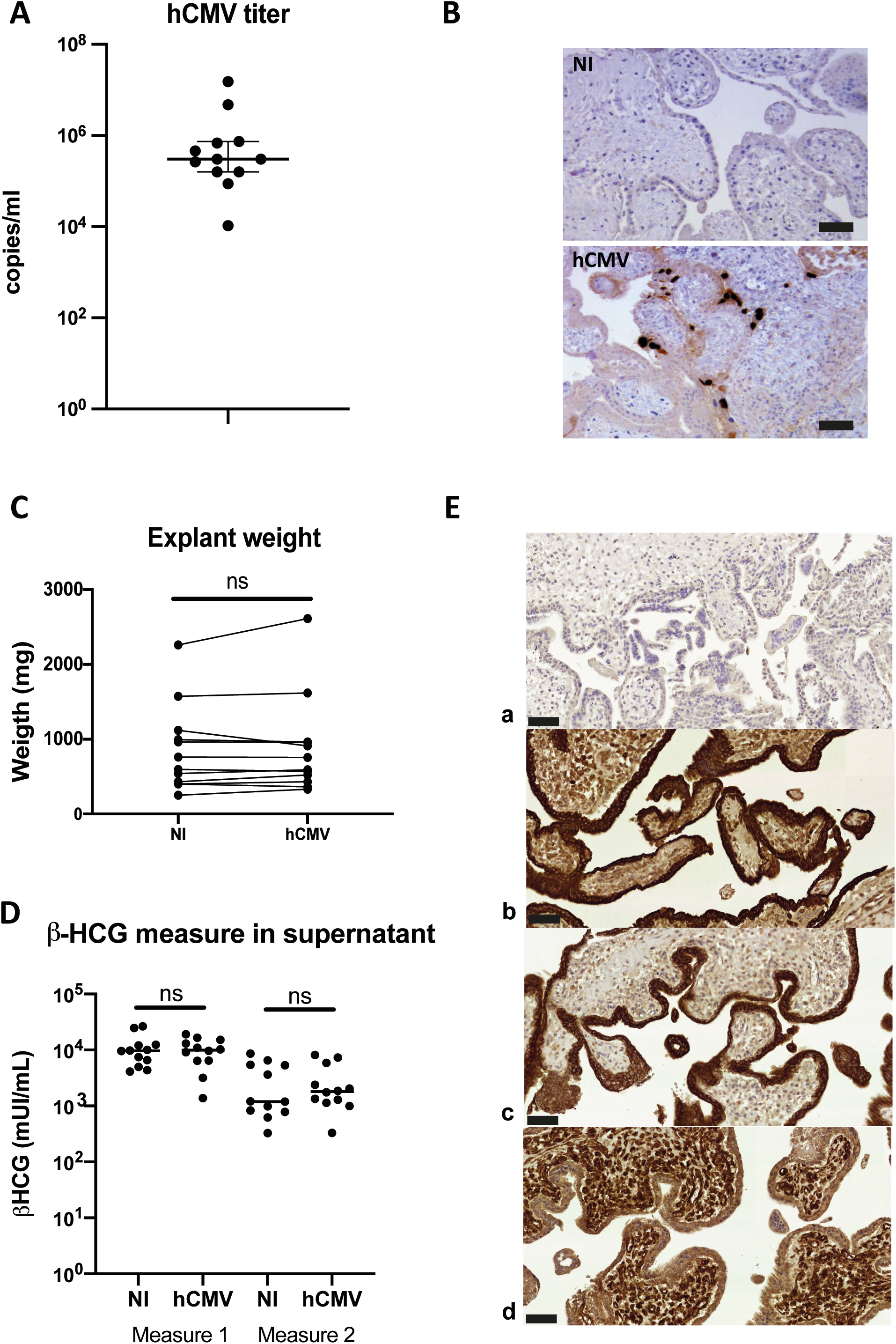
Impact of hCMV infection on placental histocultures. A) Titration of hCMV genome copies released in histoculture supernatant at day 11. On the graph is indicated the median with 95% CI. n=12 independent experiments. B) Representative examples of immuno-histochemistry performed against hCMV IE antigen at day 14 on sections of placental villi, counterstained with hematoxylin. Scale bar = 50 μm. C) Comparison of the explant weight at the end-point of the placental histocultures, performed for twelve independent experiments. Each of the placental explants were pooled per condition and weighted (NI = non-infected). ns, non-significant (*p=0,3804*) by Wilcoxon paired test. D) Comparison of the β-HCG secretion in explant supernatants was performed between non-infected (NI) *versus* hCMV-infected placenta, measured with the same timeline as presented in Figure 1A. ns, non-significant (*p=0.9697* for measure 1; *p=0,5693* for measure 2) by Wilcoxon paired test. n=12 independent experiments. E) Cross sections of immunohistochemistry and hematoxylin staining of hCMV infected placental villi from histoculture at day 14, observed by bright field microscope. a-Isotype control; b-Cytokeratin 7; c-PLAP; d-Vimentin. Image representative from at least three independent experiments. Scale bar = 50 μm.

Finally, we compared the characteristics of the sEVs secreted by non-infected or infected placental histocultures. The protocol for sEV preparation, which combines differential ultracentrifugation and gradient ultracentrifugation steps, guaranteed that viral particles did not contaminate sEV preparations, consistent with previous findings [40]. Indeed, hCMV particles are bigger and denser than sEVs and are not co-purified with sEVs upon density gradient ultracentrifugation [40]. The absence of infectious viral particles in sEV fractions was actually confirmed, by applying sEVs purified from infected placental explants to MCR5 cells and performing an anti-IE immunofluorescence assay. As expected, no IE expression was detected (Supplementary Figure 3). Moreover, no viral particle was detected by TEM in sEV preparations in all the wide field pictures examined (exemplified in Supplementary Figure 4).

When comparing yields of purified sEVs in non-infected *versus* infected histocultures, we did not observe any significant difference (Figure 7A), indicating that hCMV infection did not affect the global production of sEV by placental tissue. By TEM, sEVs prepared from infected explants displayed the same morphology (Figure 7C and Supplementary Figure 4) and relative size distribution than sEVs isolated from non-infected histocultures (Figure 7D), with no significant difference in their mean size (Figure 7E). sEVs prepared from hCMV-infected explants also expressed CD63, that was detected both by western blot (Figure 7B) and by IEM (Figure 7F-G and Supplementary Figure 5), being expressed on nearly 60% of the vesicles (Figure 7H).

**Figure 7:**
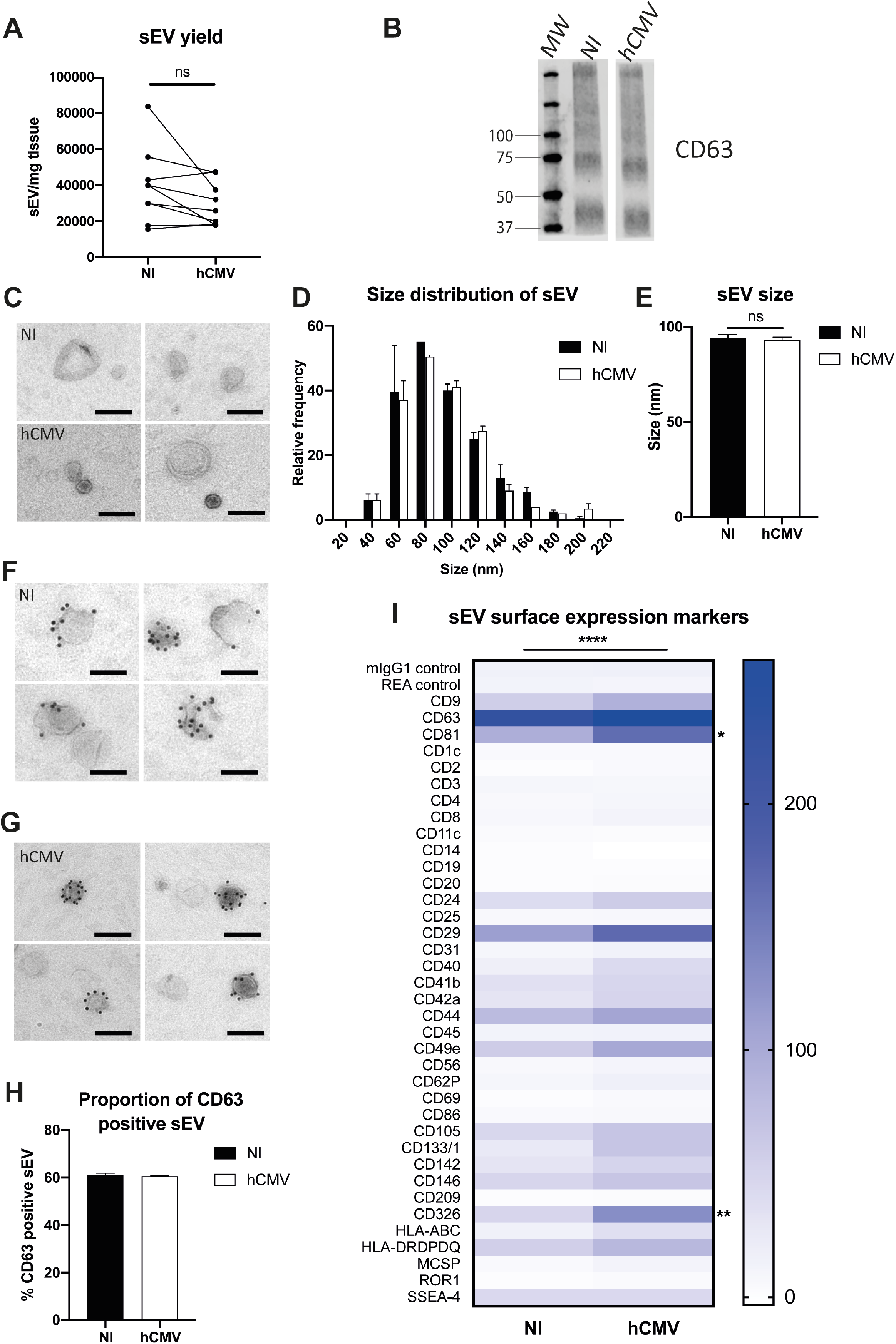
Placental sEV isolation and characterization upon hCMV infection. A) Comparison of the yield of sEV prepared per mg of placental tissue upon sEV preparation between non-infected (NI) *versus* hCMV-infected placental explants. Yield is expressed in sEV number/mg tissue, obtained from nine independent experiments. ns, non-significant by Wilcoxon paired test. B) Western blot analysis against CD63 in sEVs prepared from non-infected (NI) or infected (hCMV) placental explants. This result is representative for at least three independent experiments. CD63 appears as a smear since the non-reducing conditions of the western blot preserves its rich glycosylated pattern. MW = molecular weight. C) Placental sEV isolated from non-infected (NI) *versus* infected (hCMV) placental explants, obtained by TEM. These pictures are representative of two independent experiments. Scale bar = 100 nm. D) Frequency distribution analysis of placental sEV size, compared between non-infected (NI) *versus* infected (hCMV) placental explants. Each bar of the histogram represents the mean ± SEM of the relative frequency per bin (bin width = 20 nm) of two independent experiments. Placental sEV size were measured manually with iTEM measure tool. Total sEV count was 172 and 208 for the NI replicates; 176 and 185 for the hCMV replicates. ns, non-significant (*p=0.8814*) by nested *t*-test. E) Mean size ± SEM of placental sEV purified from non-infected (NI) or infected (hCMV) placental explants, calculated from the experiments presented in (D). ns, non-significant (*p=0.8871*) by nested *t*-test. F and G) Representative pictures of IEM analysis of placental sEV using antibodies against CD63 (gold bead size = 10 nm), purified from non-infected (NI; F) *versus* infected (hCMV; G) placental explants (n = 2). Scale bar = 100 nm. H) Percentage of placental sEV positive for CD63 (at least 1 bead counted per sEV) for two independent experiments. Total sEV count was 189 and 263 for sEV isolated from non-infected placental explants; 143 and 594 for sEV isolated from infected explants. ns, non-significant by nested *t*-test. I) Surface expression level of different proteins of sEV isolated from non-infected (NI) *versus* infected (hCMV) placental explants, based on the multiplex flow cytometry MACSPlex exosome kit assay. Results are represented by a heat-map, calculated from 3 independent experiments for different sEV markers indicated on the left column. Blue intensity is proportional to the level expression calculated in Median Fluorescence Intensity, indicated on the right of the heat-map. *p <0.0001* by 2-way ANOVA for “Infection” factor. Bonferroni post-hoc comparison test indicated significant increase for CD81 (*\*, p=0.0223*) and CD326 (*\*\*, p=0.0029*) for hCMV compared to non-infected (NI) conditions.

Finally, the surface expression levels of several proteins expressed by sEVs were examined by multiplex bead-based flow cytometry assay in both conditions (Figure 7I). To perform this assay, quantification of the sEVs by PKH67-based flow cytometry could not be realized, since it would interfere with the assay. Thus, sEV quantity was normalized between non-infected and infected conditions based on the weight of the explants, since the hCMV infection did not modify the yield of sEV secretion (Figure 7A). sEVs isolated from hCMV infected explants expressed the same markers than sEVs isolated from non-infected explants, albeit with significant differences for some of them in their expression levels upon infection (Figure 7I). A 2-way ANOVA statistical test confirmed that the infection modified the global pattern of expression of sEV surface proteins (*p<0.0001* for the “Infection” factor, no interaction with “Marker” factor). Most of the surface markers expressed in sEV isolated from infected explants showed an increased expression upon infection. Bonferroni’s multiple comparison test indicated that two markers were significantly increased: CD81 (*p=0,0223*) and CD326 (EpCAM; *p=0,0029*). In conclusion, our results indicate that hCMV infection of placental explants preserves the global secretion of sEVs that conserve the typical characteristics of exosomes, with an increase in the expression levels of some surface proteins that may be candidates as sEV markers of hCMV infection.

## DISCUSSION

An increasing number of works are centered on placental EVs and their role in physiological and pathological pregnancy. For these studies, many models have been used as a source of EVs, both *in vivo*, *ex vivo* or *in vitro*. *In vivo*, study of placental EVs isolated from blood is hard to interpret because they come from multiple tissues. The use of placental primary cells or cell lines *in vitro* is very informative but may lack important aspects of (patho)physiology occurring in a complex tissue architecture, especially during viral infection. Here, we adapted a previously established model of first trimester placental explants, which can be maintained in culture at the air/liquid interface and is permissive for hCMV replication [31, 32]. This model has also been used for different types of tissues [22], including placenta [23, 55, 56] and allows the maintenance of the tissue in culture for several days. Here, we confirmed that the integrity of the placental explants was preserved, consistent with our previous results [31, 32], with an expected pattern of β-HCG secretion along time, indicative of tissue viability [45]. Moreover, immunohistochemical analyses further established that the complex cytoarchitecture of the trophoblastic villi was well preserved at the end of the culture, even upon hCMV infection.

From these placental explants, we developed robust and reproducible conditions for the recovery and isolation of sEVs (EV-METRIC score of 100% [41]), using a combination of successive differential and density gradient ultracentrifugation steps, adapted from [38–40] and in strict accordance with MISEV guidelines [9]. We unambiguously demonstrated that our sEV preparations were pure and devoid of contaminants, as evidenced by the assessment of multiple parameters. Notably, we showed that sEV preparations presented many features of endosomal-derived exosomes, including membranous vesicles as observed by TEM, an average relative diameter around 95 nm and the presence of exosome components including CD63, CD9 and CD81. Since the subcellular origin of these vesicles cannot, however, be definitively ascertained, we have therefore chosen to keep the designation of sEVs in this manuscript [9].

Contrasting with studies where EVs are isolated from a single cell type, we examined here the global population of sEVs secreted from trophoblastic villi. The origin of the vesicles is therefore varied, reflecting those of the placental environment and enabling to assess the overall changes of the vesicles following stress. As the tissue architecture was well preserved, it is likely that the outer layers of cyto- and syncitiotrophoblasts contributed to sEV secretion. Indeed, proteins expressed by trophoblasts were actually detected on the sEV surface, including CD326, CD24 or CD49e [47, 48, 51]. We also observed the presence of proteins described for mesenchymal stem cells, like CD29, CD44 and SSEA-4, indicating that such cells also probably participate to the secretion of sEV in our model [53, 57]. Of note, even if sEV have different cellular origins, the pattern of expression of surface markers is very reproducible among sEV preparations.

We also sought to examine the impact of hCMV infection on sEVs secreted by the placental villi, because we reasoned that analysis of sEVs from first trimester placenta may be particularly well suited considering the pathophysiology of hCMV congenital infections. Indeed, hCMV efficiently disseminates from the mother to the fetus *via* an active replication in the placenta tissue [58, 59]. Consistent with previous works [31, 32, 56, 60], hCMV disseminated well in the placental explants and was released into the medium. Based on immunohistochemistry data, tissue infection levels were similar to what can be observed on placentas during natural infection [61]. Under our conditions of infection, the placental explants kept the same weight and histological structure, and continued to secrete sEVs at yields comparable to the uninfected explants.

To maximize the recovery of sEVs for a deep characterization and downstream analyses, the histoculture supernatants were pooled along the culture. Although this may either hide fluctuations and/or attenuate transient or late trends induced by the virus, we observed significant changes in the signature of sEV surface markers upon infection. A significant surface expression increase was observed notably for two proteins: CD326 and CD81. Of note, CD326 (EpCAM) has been suggested to play a role in placental development [51, 62], whereas CD81 has been recently described to play a role in hCMV entry [63, 64], although not yet for placental cells. Hence, it is tempting to speculate that the secretion of sEVs with this specific pattern upon hCMV infection may have a functional role, by influencing viral dissemination into the tissue and/or contributing to placental defects.

Currently, there is a growing interest for the search for biomarkers reflecting the state of the placenta, even of the fetus, within the placental sEVs [14, 16-18]. However, in numerous models of placental explants described to date, sEVs are generally prepared within the first 16 to 48 hours of culture, a duration that does not allow to evaluate the long-term effects of chronic stress. Hence, our model of early placental explants that can be cultured over several days appears as a very valuable tool to evaluate the impact of chronic environmental stress, including viral infection but also hypoxia or endocrine disruptors, on the secretion and composition of sEVs secreted by the placenta in early pregnancy. Ultimately, it could also open new perspectives in the search for biomarkers.

## ACKNOWLEDGMENTS

We thank the medical and paramedical staff of the gynecology unit at Paule de Viguier Hospital, who allowed us to have access to the samples, as well as the patients who agreed to participate in the study. We also warmly thank Louis Bujan and Mélanie Aubry, from the Germethèque, for their help and professionalism, and Christian Sinzger who kindly provided us with the hCMV strain and gave us precious advice for its production. We greatly thank Benjamin Rauwel, Maryse Romao, Catherine Mengelle, Francine Chauvrier, Philippe Verdy, Nicolas Kopf and Matthieu Barbet, as well as the whole ViNeDys team, for their technical assistance and their numerous advice and discussions which allowed the progress of this work. Our thanks also go to Anne-Laure Iscache, Valérie Duplan-Eche and Fatima L’Faqihi-Olive, from the cytometry facility of the Center for Pathophysiology of Toulouse Purpan, as well as to Florence Capilla and Annie Alloy, from the histology facility Gentoul Anexplo. We also thank Laurence Nieto and Evert Haanappel from the Institute of Pharmacology and Structural Biology, who allowed us to have access to the Nanosight device, and for their many advices.

This project has received financial support from the French Society of Neonatology, the French Biomedicine Agency, the Réseau Mère-Enfant de la Francophonie. Our work was also financially supported by institutional grants from Inserm, CNRS and Toulouse 3 University. This project is part of the doctorate thesis of Mathilde Bergamelli, who was funded by the Ministry of Education and Research (MESR). The TEM experiments were performed on PICT-IBiSA, Institut Curie, Paris, member of the France-BioImaging national research infrastructure, and were supported by the French National Research Agency through the “Investments for the Future” program (France-BioImaging, ANR-11-INSB-04), supported by the CelTisPhyBio Labex (N° ANR-11-LB0038) part of the IDEX PSL (N°ANR-10-IDEX-0001-02 PSL).

## DISCLOSURE STATEMENT

The authors report no conflict of interest.

## Notes

### Competing Interest Statement

The authors have declared no competing interest.

