## Supplementary material for "Human cytomegalovirus infection changes the pattern of surface markers of small extracellular vesicles isolated from first trimester placental histocultures": supp Table 1

**Supplementary Table 1: Comparison of EV counting by flow cytometry and NTA methods**

Unit: particle/mL

| Experiment n° | flow cytometry | NTA |
|---|---|---|
| #37 | 1,45E+08 | 1,55E+08 |
| #38 | 8,80E+07 | 4,08E+08 |
| #39 | 1,00E+08 | 1,66E+08 |
| #40 | 3,00E+08 | 2,63E+08 |
