## supplementary Figures for "Human cytomegalovirus infection changes the pattern of surface markers of small extracellular vesicles isolated from first trimester placental histocultures"

Electron microscopy of small EVs isolated from non-infected placental histoculture

15000x

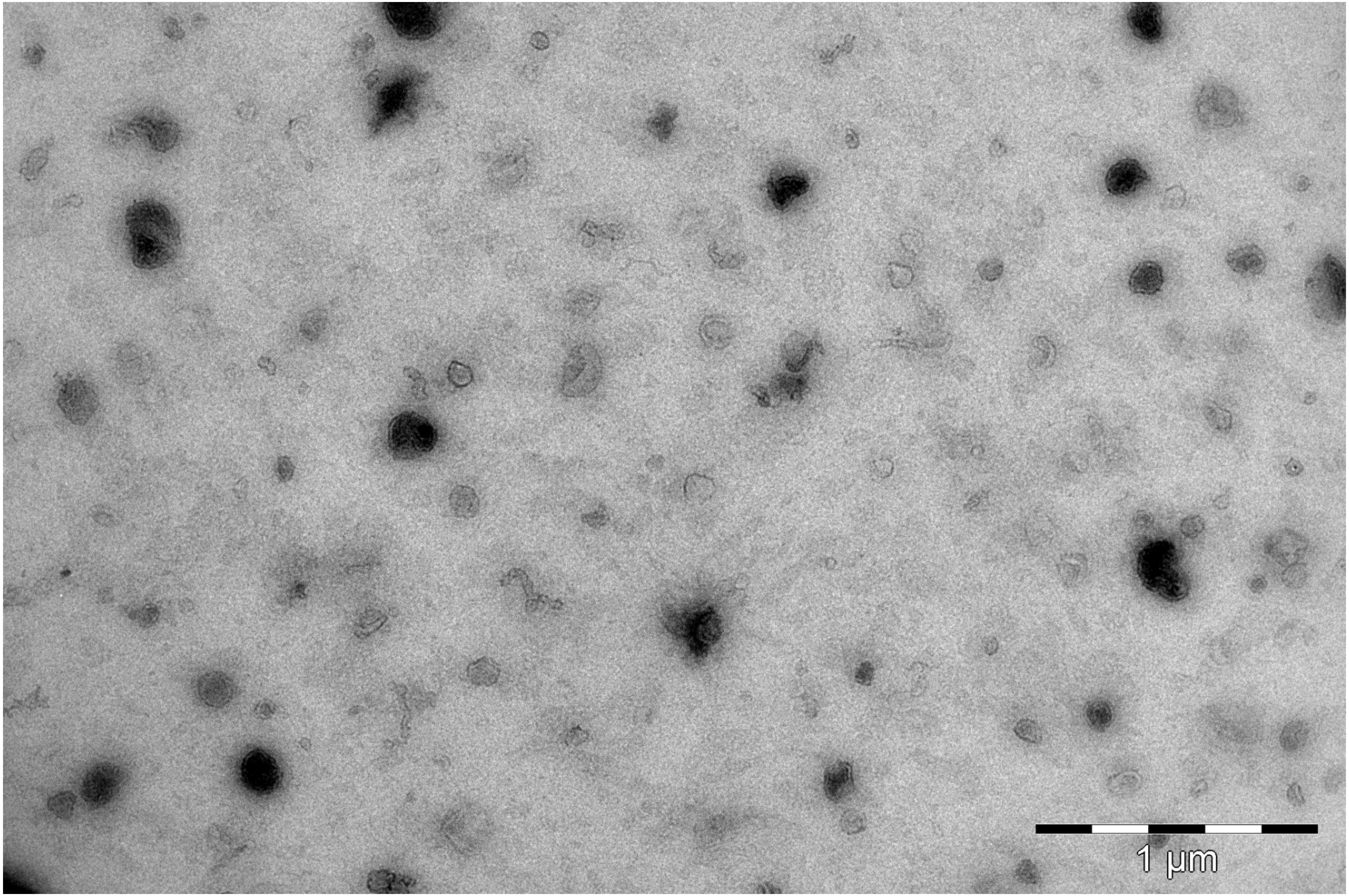

Immuno electron microscopy anti-CD63 of small EVs isolated from non-infected placental histoculture 15000x

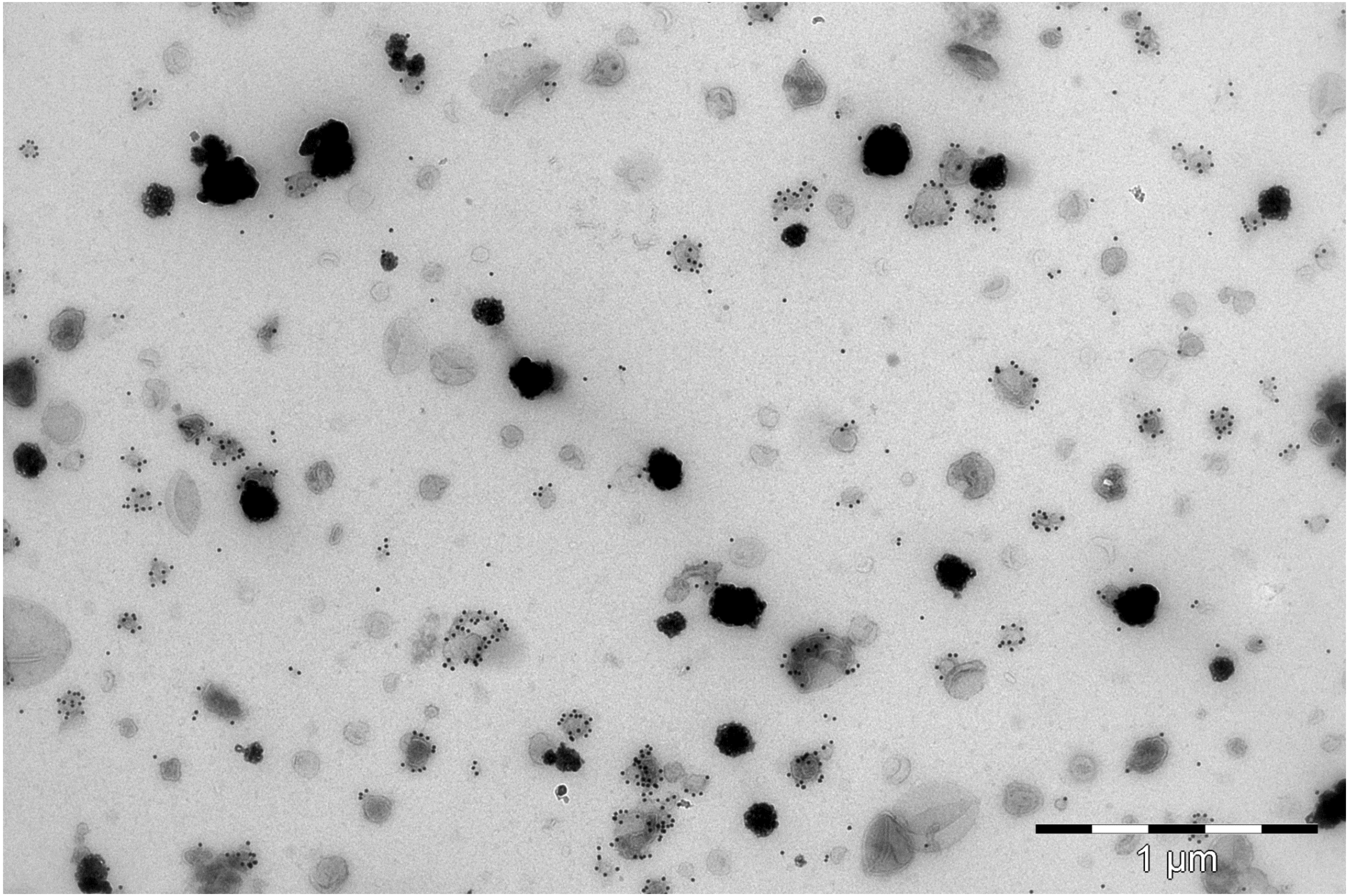

Placental small EVs preparation are devoid of infectious virus. Small EV preparations (left panel) or hCMV at a multiplicity of infection of 10 (right panel) were incubated with MCR5 cells during 24 h. Immunofluorescence was then performed against viral IE antigen (green: IE; blue: DAPI). Magnification = 20 X. The low density of the cells on the right panel is due to the high mortality rate consequent to the virus infection.

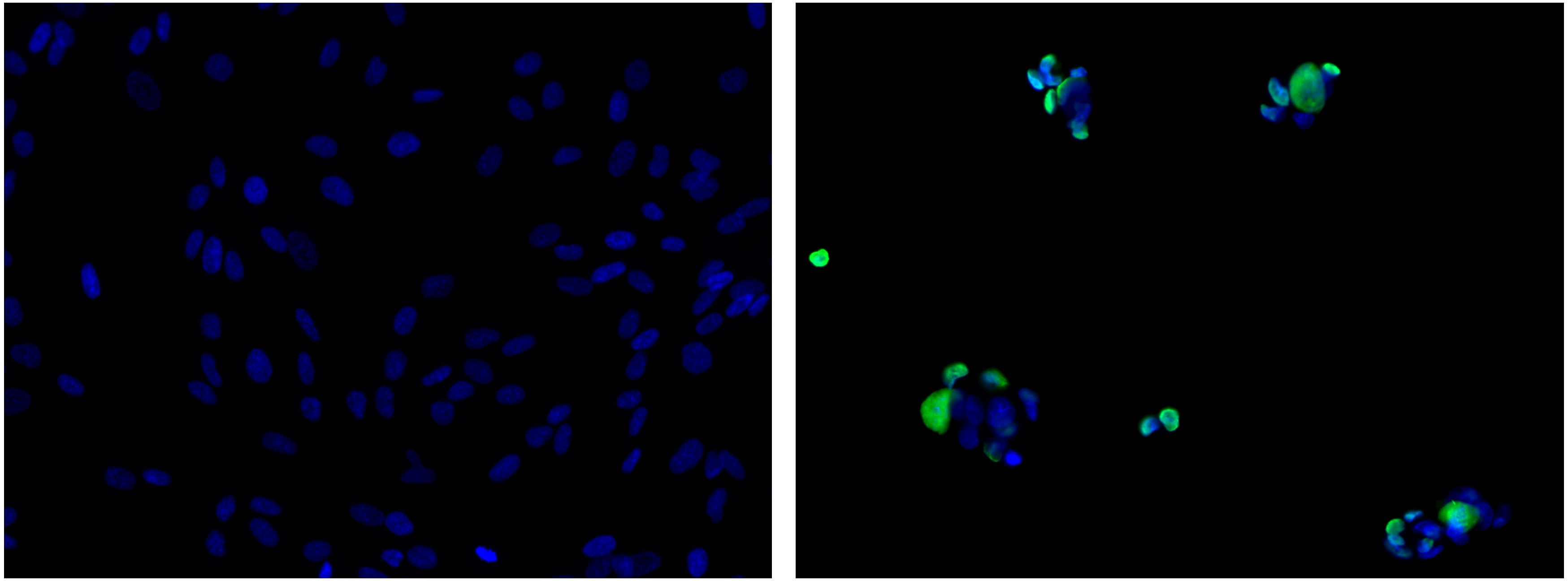

Electron microscopy of small EVs isolated from hCMV-infected placental histoculture
15000x

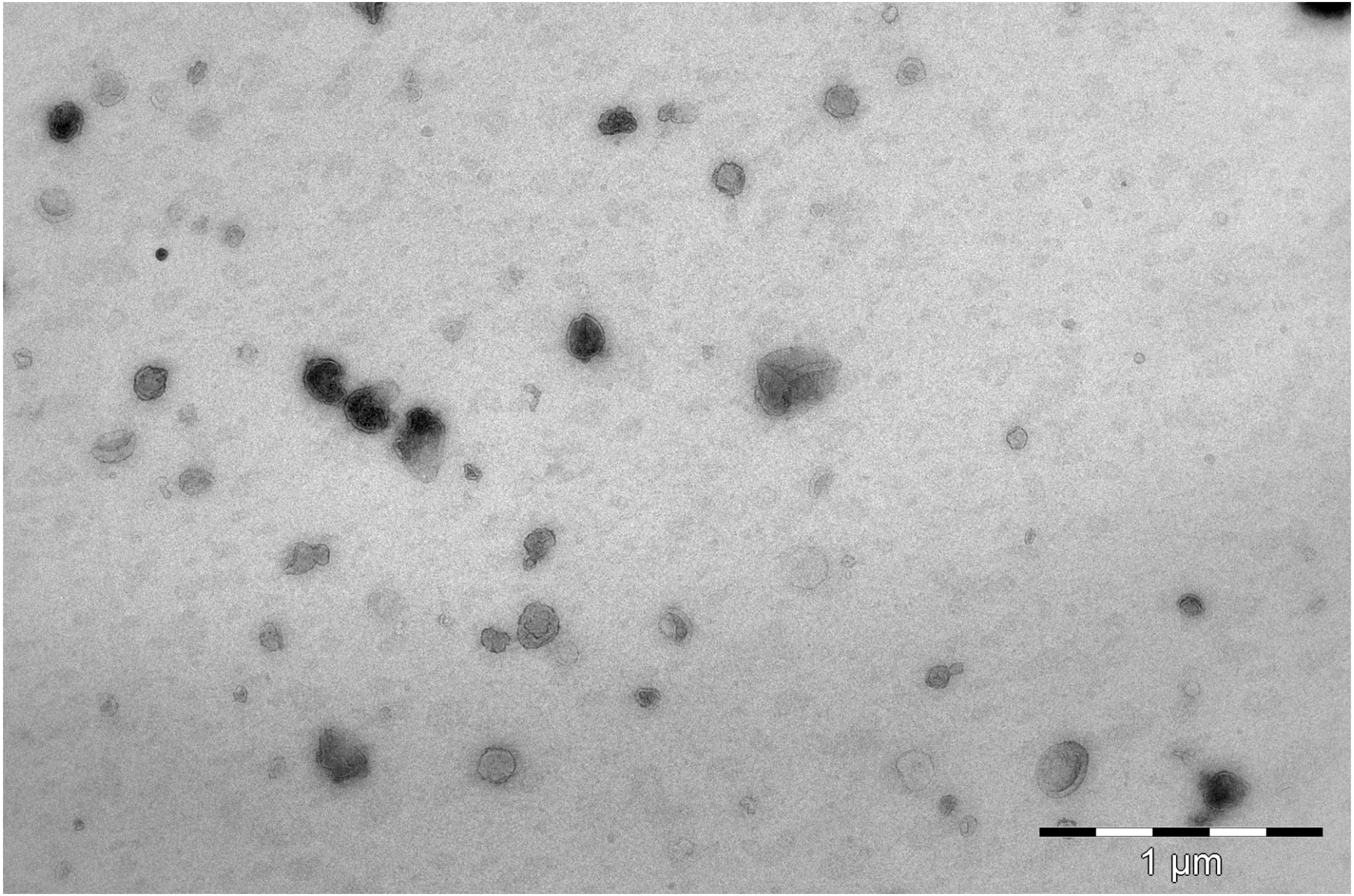

Immuno electron microscopy anti-CD63 of small EVs isolated from hCMV-infected placental histoculture
15000x

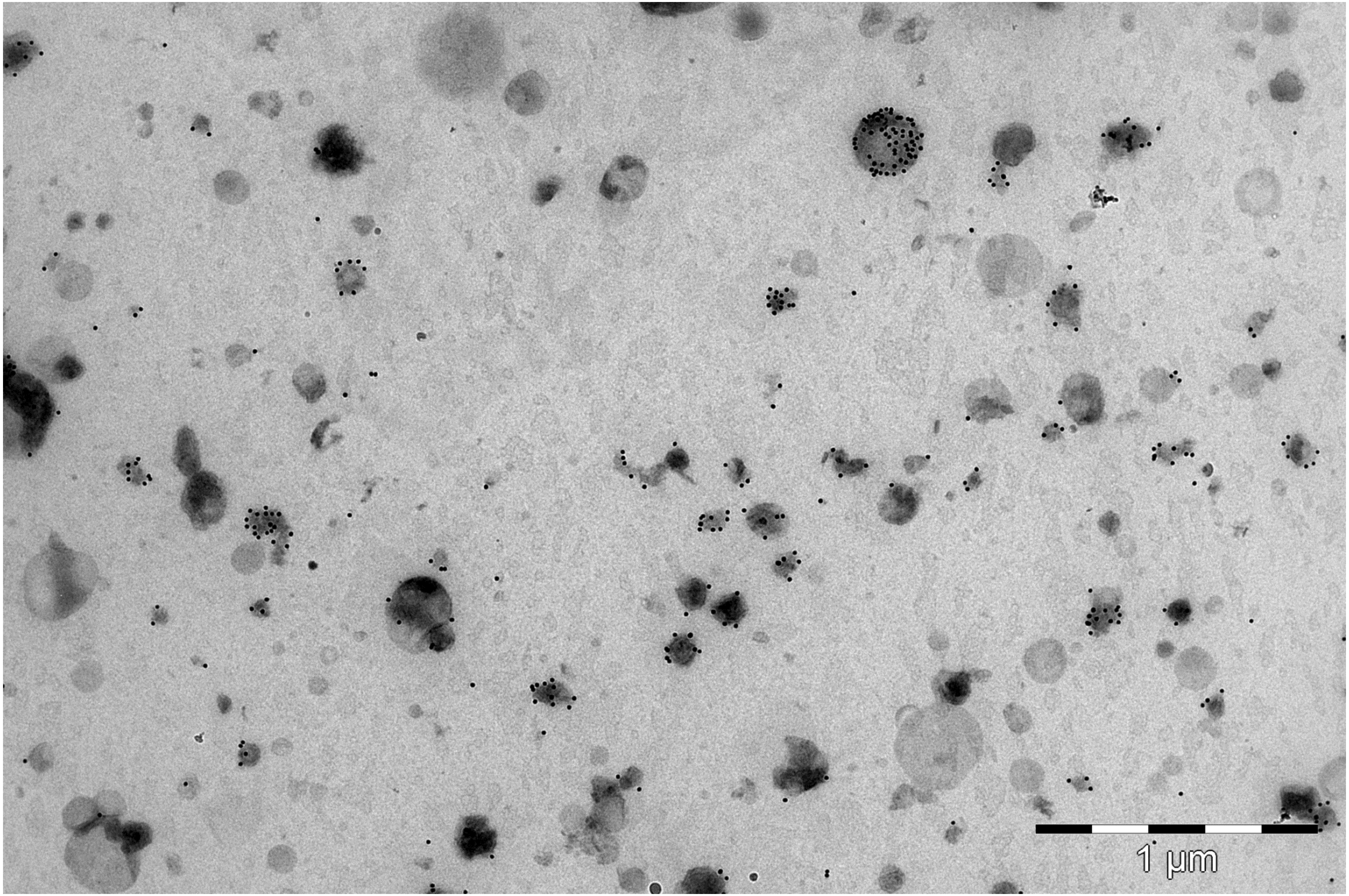
